# A CT-to-brain-atlas mapping framework using cranial base landmarks in marmosets

**DOI:** 10.64898/2026.11.18.767736

**Authors:** Haruya Yagishita, Masanori Kawabata, Yuji Nagai, Yuki Hori, Yuka Hirayama, Teppei Ebina, Keisuke Sehara, Kei Oyama, Masanori Matsuzaki, Takafumi Minamimoto, Yoshikazu Isomura

## Abstract

**Objective:** The common marmoset (*Callithrix jacchus*), a small, experimentally tractable nonhuman primate, is increasingly used as a model for studying higher brain functions. However, substantial individual differences in its skull and brain limit the accuracy of stereotaxic coordinates. Magnetic resonance imaging (MRI) of each animal improves accuracy but is costly. Computed tomography (CT) is easy to acquire and shows the skull and implanted devices, but not the brain. We therefore aimed to map CT images to a brain atlas without MRI of each animal.

**Approach:** Six marmosets with both CT and MR images served as references. A target CT image was registered to a reference CT image and mapped to the atlas via the reference’s MR image. For this CT-to-CT registration, we compared intensity-based registration with landmark-based registration using eight cranial base landmarks. We also averaged mappings across multiple references. Mapping error was measured against the mapping via the target’s own MR image.

**Main results:** With a single reference, landmark-based registration had a significantly smaller error than intensity-based registration. Averaging the mappings from five references improved the accuracy (0.45 mm), and five or more landmarks gave almost the same accuracy as all eight. Landmark-based registration also worked better on CT images from a different scanner and on postoperative CT images with implants (three and two marmosets, respectively). Using our new open-source software, Marmoset Atlas Mapper, we assigned Neuropixels recording sites to brain regions on postoperative CT, and the region boundaries were qualitatively consistent with boundaries in local field potential correlations.

**Significance:** Cranial base landmarks and multiple references allow CT-to-atlas mapping with a submillimeter mean error without MRI of each animal, even with image artifacts or partly missing landmarks. This enables *in vivo* localization of implanted devices, including those in deep structures, facilitating studies of higher brain functions in marmosets.

## Introduction

The common marmoset (*Callithrix jacchus*) has been increasingly used as a nonhuman primate model in neuroscience research owing to its small body size, experimental tractability, relatively short generation time, and amenability to genetic modification [1–4]. More recently, electrophysiological recordings with multichannel silicon probes, including high-density Neuropixels [5–8], and optogenetic manipulations [9–13] have been performed in marmosets. Interpreting the results of such experiments requires accurate targeting of the intended brain regions and verification of the recording and manipulation sites. In particular, localization with submillimeter accuracy is important for distinguishing adjacent regions in the small marmoset brain [14].

In neuroscience studies using marmosets, stereotaxic surgery, in which targets are specified by coordinates relative to cranial landmarks as in standard brain atlases [15,16], is commonly used to insert electrodes [5] and to inject tracers [17,18] or viral vectors [13,19]. As in other species, the recording and injection sites are generally verified by postmortem histological analysis [20]. However, this approach has three limitations. First, marmosets show large interindividual variability in the shape and size of the skull and brain [14,21], and the position of bregma relative to the brain varies with a standard deviation of approximately 1 mm along the anteroposterior axis [14]. Consequently, it is difficult to reach targets with submillimeter accuracy on the basis of stereotaxic coordinates alone. In addition, because the marmoset cortex is largely lissencephalic, it provides few reliable sulcal landmarks for localizing cortical areas from the brain surface [14]. Second, because histological verification is possible only after the experiment has ended, whether the target has been reached cannot be confirmed during the experiment [22]. Third, when insertions are repeated in the same animal, it is difficult to match each track identified in histological sections to the corresponding recording session.

Over the past decade, standard templates and brain atlases based on magnetic resonance imaging (MRI) have been developed for the marmoset, including the Brain/MINDS 3D Digital Marmoset Brain Atlas [23], its version 2.0 (BMA 2.0) [24], and the Marmoset Brain Mapping atlases [21,25–28]. MRI-guided stereotaxic surgery using subject-specific MR images has also been reported in marmosets [14,29]. These approaches enable targets to be planned on the basis of each animal’s MR images and the positions of inserted devices to be verified by integration with postoperative images, and can therefore address the limitations described above. However, acquiring high-resolution MR images of each animal requires access to appropriate imaging facilities, as well as time and costs for imaging and animal management. Consequently, the need for subject-specific MRI limits a key advantage of the marmoset model, namely, the ease with which relatively large numbers of animals can be studied.

Computed tomography (CT), on the other hand, is faster and simpler than MRI and clearly visualizes the skull and implanted devices, although it provides poor soft-tissue contrast in the brain. In rodents, which show relatively small interindividual variability in brain structure, several methods have been developed to localize implanted electrodes and optic fibers by registering CT images to brain atlases or histological templates, and their accuracy has been validated against histology [22,30,31]. By contrast, in primates, which show greater interindividual variability in brain size and shape, CT has typically been co-registered with MR images to achieve accurate localization [32]. In marmosets, CT has been used to design recording chambers and to inspect implants postoperatively, and target and recording-site locations have been estimated by fitting an MRI-based template brain to the skull on CT [7,12]. However, these studies did not quantify localization errors relative to MRI or histology. Thus, given the large interindividual variability in marmosets, it remains unclear how accurately CT images can be mapped to a brain atlas and which parts of the skull should be used for registration.

To address these questions, we constructed a registration pathway that maps the CT coordinates of a target marmoset into the BMA 2.0 space via a reference marmoset for which both CT and MR images are available. For CT-to-CT registration between the target and reference marmosets, we compared intensity-based registration, which aligns whole images on the basis of signal intensity, with landmark-based registration using landmarks located mainly on the cranial base or including those on the calvaria. We also examined whether averaging the mappings obtained via multiple reference marmosets reduces mapping error. Mapping error was quantified relative to the mapping obtained through the target marmoset’s own MR image. We show that combining cranial base landmarks with multiple reference marmosets allows CT images to be mapped to the atlas with a submillimeter mean error without requiring MRI of the target marmoset. We further show that the pathway can be applied to CT images acquired with a different scanner. Moreover, because cranial base landmarks remain identifiable even when implants are mounted on the calvaria, the pathway can also be applied to postoperative CT images, allowing implanted devices to be localized in atlas space. Finally, we implemented the pathway in Marmoset Atlas Mapper, a new open-source software package. As an application, we used this software to estimate the atlas-defined brain regions of Neuropixels recording sites.

## Materials and Methods

### Experimental animals

All experiments were performed with the approval of the Animal Experiment Committees of the Institute of Science Tokyo (approval numbers: A2024-163A, A2025-070C, and A2026-040A), the Animal Ethics Committee of the National Institutes for Quantum Science and Technology (QST; approval number: 11-1038), and the Institutional Animal Care and Use Committee of the University of Tokyo (UT; approval numbers: P20-079 and A2023M018-08). All procedures followed the Guide for the Care and Use of Nonhuman Primates in Neuroscience Research (The Japan Neuroscience Society; https://www.jnss.org/en/animal_primates). This study is reported in accordance with the ARRIVE guidelines (Animal Research: Reporting of In Vivo Experiments).

A total of 11 common marmosets (*Callithrix jacchus*; 7 males and 4 females; age, 1–11 years; body weight, 278–482 g) were used. *In vivo* CT and MR images were acquired from each of nine marmosets. Six of these marmosets were scanned at QST and used as the evaluation dataset, and the other three were scanned at UT with a higher-resolution CT scanner and used as the external dataset. The remaining two marmosets underwent Neuropixels probe implantation and were used for electrophysiological recordings and postoperative CT scanning at UT (three scans in total). Details of the individual animals are summarized in Table 1.

**Table 1.** Information on the marmosets used in this study. The columns show the dataset, animal ID, sex, imaging facility, and age and body weight at the CT and MRI scans of each marmoset. UT5 underwent two postoperative CT scans; body weights at the first and second scans are shown. QST, National Institutes for Quantum Science and Technology; UT, University of Tokyo; N/A, not acquired.

| **Dataset** | **Animal ID** | **Sex** | **Imaging facility** | **Age at scan (years)** |  | **Body weight at scan (g)** |  |
|---|---|---|---|---|---|---|---|
|  |  |  |  | **CT** | **MRI** | **CT** | **MRI** |
| Evaluation | HH0312F | Female | QST | 11 | 11 | 278 | 278 |
| Evaluation | HH0424M | Male | QST | 2 | 1 | 316 | 310 |
| Evaluation | HH0428M | Male | QST | 8 | 8 | 344 | 344 |
| Evaluation | HJ0285F | Female | QST | 11 | 11 | 338 | 356 |
| Evaluation | J0476M | Male | QST | 5 | 5 | 361 | 360 |
| Evaluation | J0477M | Male | QST | 2 | 2 | 411 | 411 |
| External | UT1 | Female | UT | 3 | 3 | 380 | 387 |
| External | UT2 | Male | UT | 2 | 2 | 362 | 364 |
| External | UT3 | Female | UT | 2 | 2 | 414 | 428 |
| Electrophysiology | UT4 | Male | UT | 5 | N/A | 482 | N/A |
| Electrophysiology | UT5 | Male | UT | 4 | N/A | 397, 356 | N/A |

### CT and MR image acquisition

In the evaluation dataset, CT and MR images were acquired at QST. CT images were acquired under anesthesia with ketamine (25 mg/kg, i.m.) and xylazine (2 mg/kg, i.m.) using a cone-beam CT system (Accuitomo 170; J. Morita, Osaka, Japan) with a tube voltage of 90 kVp, a tube current of 5 mA, an exposure time of 17.5 s, a field of view (FOV) of 140 mm (diameter) × 100 mm (height), and a voxel size of 0.25 × 0.25 × 0.5 mm. T2-weighted MR images were acquired under 2.0–2.5% isoflurane anesthesia with spontaneous respiration using a 7-T scanner (BioSpec 70/20; Bruker BioSpin, Ettlingen, Germany) with a rapid acquisition with relaxation enhancement (RARE) sequence (repetition time, 6,000 ms; echo time, 22.18 ms; RARE factor, 8; FOV, 47.7 × 47.7 mm; matrix, 191 × 191; voxel size, 0.25 × 0.25 × 0.25 mm; 12 averages). In the external dataset, CT and MR images were acquired at UT. CT images were acquired under anesthesia with ketamine (15 mg/kg, i.m.) and xylazine (0.75 mg/kg, i.m.) using a micro-CT scanner (SkyScan 1276; Bruker, Kontich, Belgium) with a tube voltage of 60 kV, a tube current of 40 µA, an Al + Cu filter, an exposure time of 800 ms per projection, and a rotation step of 1.8° over 360°. Each projection image consisted of 1,024 × 1,024 pixels with a pixel size of 36.556 µm. To cover the entire skull, two scans were performed for each marmoset, with a total X-ray exposure time of approximately 540 s. Three-dimensional images were reconstructed using NRecon (Bruker) with the default parameters. T2-weighted MR images were acquired under 2.5–3.5% isoflurane anesthesia with spontaneous respiration using a 3-T scanner (BioSpec 3T; Bruker BioSpin, Ettlingen, Germany) with a RARE sequence (repetition time, 4,000 ms; effective echo time, 24 ms; RARE factor, 32 for UT1 and 64 for UT2 and UT3; FOV, 40 × 40 × 30 mm; matrix, 160 × 160 × 120; voxel size, 0.25 × 0.25 × 0.25 mm; 4 averages). Postoperative CT images of the two Neuropixels-implanted marmosets were acquired with the same scanner and procedures as in the external dataset, except for a tube voltage of 50 kV, a tube current of 100 µA, a 1-mm Al filter, an exposure time of 1,600 ms per projection, and a rotation step of 1.0°.

### Registration of CT and MR images to the atlas

The population-averaged *in vivo* T2-weighted MR image of the Brain/MINDS 3D Digital Marmoset Brain Atlas version 2.0 (BMA 2.0) was used as the atlas [24]. The MR image of each marmoset was corrected for intensity inhomogeneity using N4 bias-field correction [33] with a shrink factor of 4, 50 iterations at each of four levels, and a convergence threshold of 10^−4^. After manual initial alignment, the corrected MR image was registered to the atlas with rigid, affine, and BSplineSyN transformations in this order, using the Advanced Normalization Tools (ANTs) software package [34]. The CT of each marmoset was manually aligned to the MR image of the same individual and then registered to it with a rigid transformation using ANTs. These transformations were concatenated to obtain the CT-to-atlas transformation of each marmoset. All registration results were visually inspected on superimposed images.

For linear registration, Mattes mutual information (32 bins; regular sampling, 25%) was used as the similarity metric with a gradient step of 0.1. A four-level multiresolution scheme was used, with shrink factors of 8, 4, 2, and 1, smoothing sigmas of 3, 2, 1, and 0 voxels, and maximum iterations of 1,000, 500, 250, and 100, respectively. The convergence threshold was set to 10^−6^ with a convergence window of 10 iterations. For nonlinear registration, BSplineSyN[0.1,26,0,3] was used with local cross-correlation (radius, 4 voxels) as the similarity metric and maximum iterations of 100, 70, 50, and 20.

### CT-to-CT registration

Thirteen skull landmarks (LM01–13) were manually placed on the *in vivo* CT image of each marmoset (Figure 1, Table 2), and three landmark sets were defined: Total (all 13 landmarks), Lower (eight landmarks located mainly on the cranial base, LM01– 08), and Upper (eight landmarks including those on the calvaria, LM06–13). Landmark-based registration was performed using vtkLandmarkTransform with two transformation models: rigid (rotation and translation; 6 degrees of freedom) and similarity (rotation, translation, and isotropic scaling; 7 degrees of freedom). Intensity-based registration was performed using ANTs; a rigid transformation was estimated for the rigid model, and a rigid transformation followed by a similarity transformation was estimated for the similarity model. All registrations were initialized by manual rigid alignment, and the same initial alignment was shared across registration methods for each pair of marmosets.

**Figure 1.**
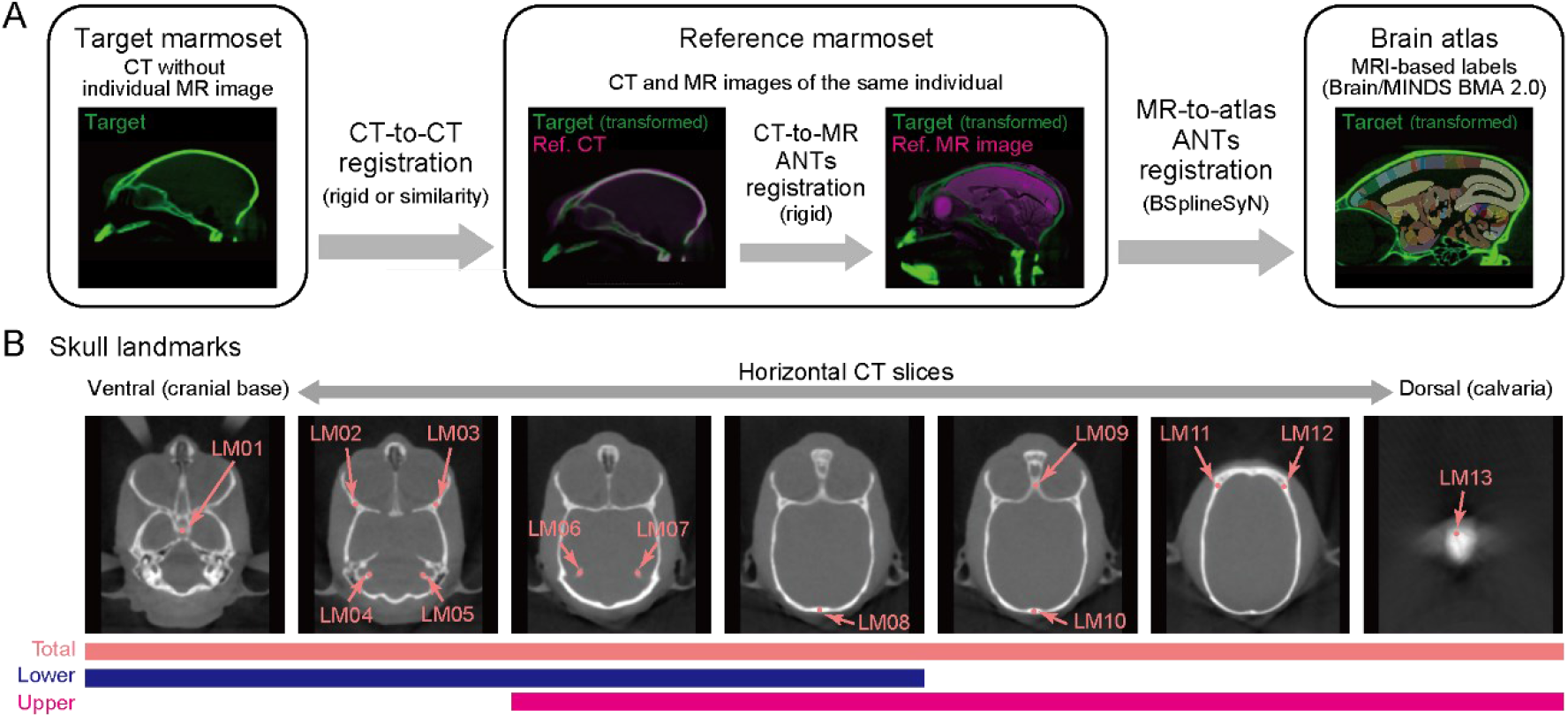
Marmoset CT-to-brain-atlas registration pipeline using a reference marmoset. (A) Schematic of the registration pipeline. A reference marmoset for which both CT and MR images were available was used to establish the registration pathway. Three transformations were concatenated to map a target CT image into the Brain/MINDS 3D Digital Marmoset Brain Atlas version 2.0 (BMA 2.0) space: landmark-based CT-to-CT registration from the target to the reference marmoset, rigid CT-to-MR registration within the reference marmoset, and nonlinear MR-to-atlas registration. CT-to-MR and MR-to-atlas registrations were performed with the Advanced Normalization Tools (ANTs) software package. Green, target CT; magenta, reference CT and MR images. (B) Skull landmarks used for landmark-based interindividual CT registration. Pink arrows indicate the positions of 13 landmarks (LM01–13) on representative CT sections. Anatomical definitions of the landmarks are provided in Table 2. The color bars show the three landmark sets used for registration and their constituent landmarks: Total (pink), comprising all 13 landmarks (LM01–13); Lower (blue), comprising eight landmarks (LM01–08); and Upper (magenta), comprising eight landmarks (LM06–13).

**Table 2.**
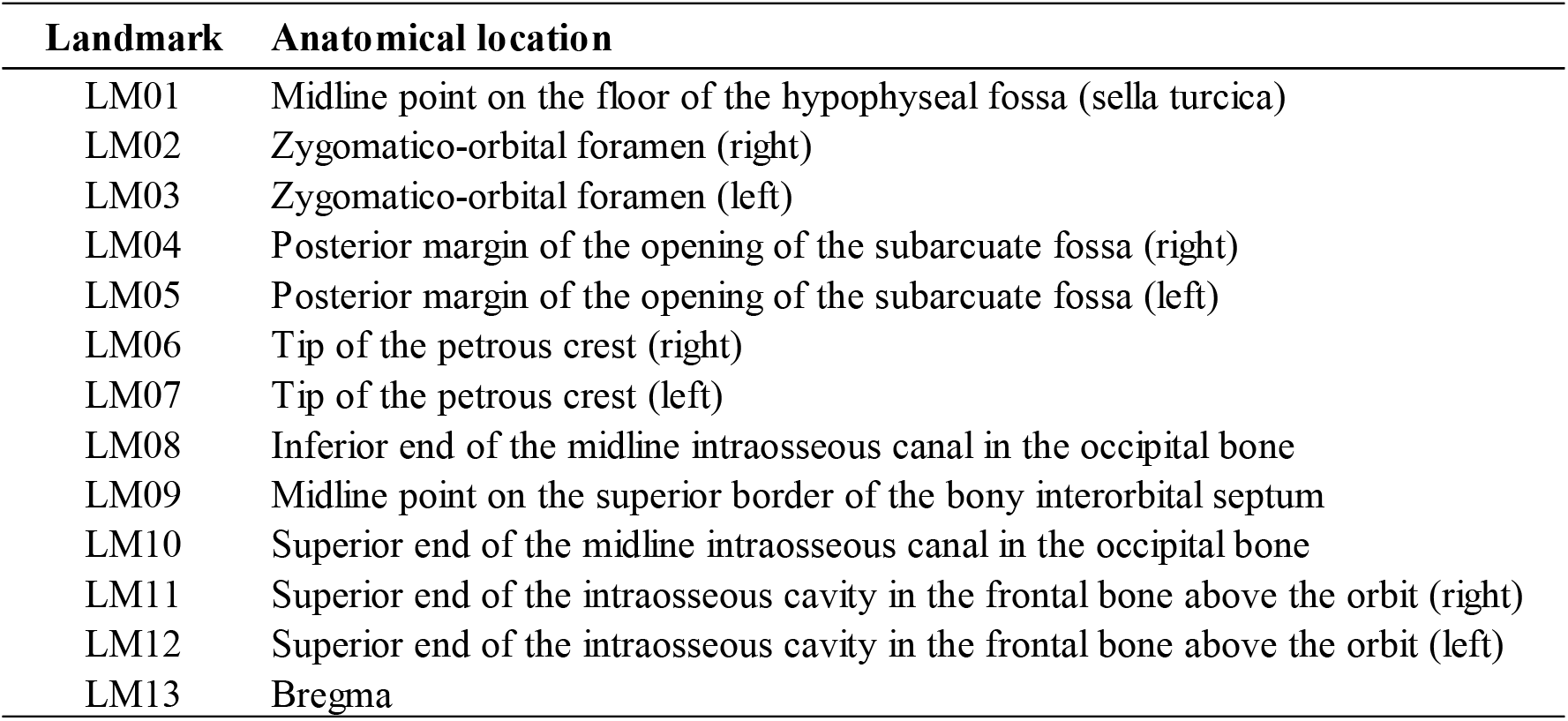
Anatomical definitions of the skull landmarks (LM01–13). The columns show the landmark number and the anatomical location of each landmark. LM02/03, LM04/05, LM06/07, and LM11/12 are bilateral pairs of left and right landmarks; the remaining landmarks are located on the midline. The Total set comprises all 13 landmarks, the Lower set comprises LM01–08, and the Upper set comprises LM06– 13.

| **Landmark** | **Anatomical location** |
|---|---|
| LM01 | Midline point on the floor of the hypophyseal fossa (sella turcica) |
| LM02 | Zygomatico-orbital foramen (right) |
| LM03 | Zygomatico-orbital foramen (left) |
| LM04 | Posterior margin of the opening of the subarcuate fossa (right) |
| LM05 | Posterior margin of the opening of the subarcuate fossa (left) |
| LM06 | Tip of the petrous crest (right) |
| LM07 | Tip of the petrous crest (left) |
| LM08 | Inferior end of the midline intraosseous canal in the occipital bone |
| LM09 | Midline point on the superior border of the bony interorbital septum |
| LM10 | Superior end of the midline intraosseous canal in the occipital bone |
| LM11 | Superior end of the intraosseous cavity in the frontal bone above the orbit (right) |
| LM12 | Superior end of the intraosseous cavity in the frontal bone above the orbit (left) |
| LM13 | Bregma |

### Multi-reference registration

The mapping from the target CT to the atlas via reference marmoset *i* was defined as follows:

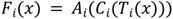

where *x* is a coordinate in the target CT, *T_i_* is the transformation from the target CT to the CT of reference marmoset *i*, and *C_i_* and *A_i_* are the CT-to-MR and MR-to-atlas transformations of reference marmoset *i*, respectively.

When *N*_ref_ reference marmosets were used, the atlas coordinates obtained via the individual references were averaged with equal weights to obtain the consensus mapping:

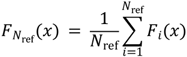

The displacements *F_i_*(*x*) − *x* obtained via the individual references were averaged on a common grid with 0.5-mm spacing covering the entire target CT with a 1-mm margin, yielding a mean displacement field; values between grid points were obtained by linear interpolation using SimpleITK. To superimpose the atlas MR image and labels onto the target CT, the atlas coordinate corresponding to each CT voxel was computed using this mean displacement field, and the atlas MR image and labels were sampled at these coordinates using linear and nearest-neighbor interpolation, respectively.

This registration pathway was implemented in a Windows desktop application, Marmoset Atlas Mapper, written in Python 3.12 with PySide6 (see Data and code availability).

### Evaluation of mapping accuracy

In the evaluation dataset, each marmoset was used in turn as the target marmoset, and the remaining five marmosets were used as candidate references. In the external dataset, each marmoset was used as the target marmoset, with all six marmosets in the evaluation dataset as references (*N*_ref_ = 6). An intracranial region was created on each target CT and visually confirmed, and a 1-mm grid was placed within this region (7,867– 9,513 and 8,490–8,940 points per target marmoset in the evaluation and external datasets, respectively). The mapping through the target marmoset’s own CT–MR–atlas pathway, *D*(*x*), was used as the within-subject mapping and served as the benchmark (Figure 2). The mapping error at each grid point was defined as the Euclidean distance between the cross-subject mapping *F_N_*_ref_(*x*) and *D*(*x*). The mean error of each target marmoset was computed by the following formula:

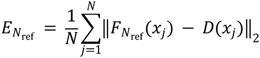

**Figure 2.**
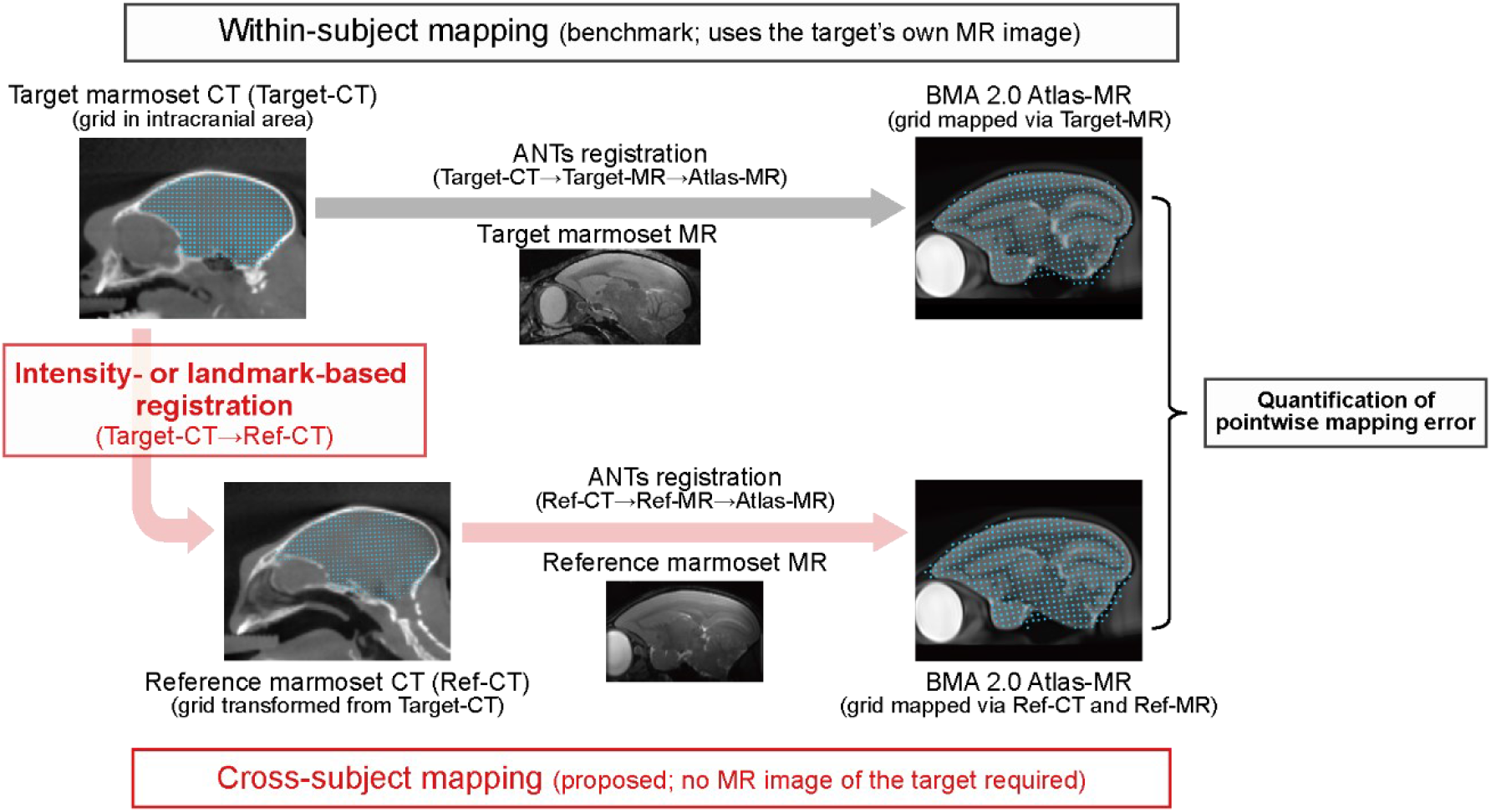
Quantification of CT-to-atlas mapping error with within-subject mapping as the benchmark. An intracranial grid of points (cyan) defined in the CT image of the target marmoset (Target-CT) was mapped to atlas space through two registration pathways. In the within-subject mapping (upper pathway, gray arrow), the points were transformed using rigid CT-to-MR registration within the target marmoset, followed by MR-to-atlas registration. In the cross-subject mapping (lower pathway, red arrows), the same points were first mapped to the CT image of the reference marmoset (Ref-CT) using intensity- or landmark-based CT-to-CT registration, then transformed into atlas space using the reference marmoset’s CT-to-MR and MR-to-atlas transformations. Pointwise mapping error was quantified as the Euclidean distance between the corresponding point coordinates obtained through the two pathways in atlas space. The target marmoset’s MR image was used to establish the within-subject mapping for validation but was not required for the cross-subject mapping. Target-MR and Ref-MR, MR images of the target and reference marmosets; Atlas-MR, atlas MR image; ANTs, Advanced Normalization Tools.

where *x_j_* is the *j*th grid point and *N* is the number of grid points. Errors along the anteroposterior (AP), dorsoventral (DV), and mediolateral (ML) axes were computed in the same manner by averaging the absolute values of the corresponding components of the error vector. For each *N*_ref_ from 1 to 5 in the evaluation dataset, errors were computed for all combinations of *N*_ref_ references selected from the five candidates and averaged within each target marmoset.

Spatial profiles of mapping errors were obtained by dividing the AP, DV, and ML positions into 1-mm bins and averaging the Euclidean errors within each bin for each target marmoset. The origin of the AP and DV axes was set at bregma (LM13) mapped to the atlas by *D*(*x*), and the origin of the ML axis was set at the midsagittal plane of the atlas (R = 0 in the RAS coordinate system), with rightward defined as positive.

### Effects of landmark number and contributions of individual landmarks

Using the eight Lower landmarks, the number of landmarks used for registration was varied from three to eight, and errors were computed for all possible combinations under the rigid model (*N*_ref_ = 5). For each number of landmarks, errors were averaged across combinations within each target marmoset and then summarized across the six target marmosets. In addition, the difference in error from the eight-landmark configuration was computed for each number of landmarks in each target marmoset, and its mean and 95% confidence interval (CI) were calculated.

The importance of each landmark *l* was defined as the decrease in the mean error when *l* was added to a combination *S* of three Lower landmarks not containing *l*, as follows:

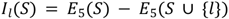

where *E*_5_(*S*) denotes the mean error obtained with combination *S* and *N*_ref_ = 5. Positive values indicate that adding the landmark reduced the error. Importance values were computed for all possible combinations *S* and averaged. For bilateral landmark pairs (LM02/03, LM04/05, and LM06/07), the importance values of the left and right landmarks were averaged.

### Reproducibility of landmark placement

Two raters with no prior experience placed the eight Lower landmarks on the CT images of the six marmosets in the evaluation dataset using Marmoset Atlas Mapper after an explanation of the landmark definitions (Table 2). The landmarks of a reference marmoset other than the target were displayed as a guide, and each landmark was placed once. The distances between the raters’ landmarks and the original landmarks placed by H. Y. were calculated, and the mapping errors were recomputed using the raters’ landmarks on the target CT (*N*_ref_ = 5).

### Stereotaxic surgery and Neuropixels probe implantation

All surgeries were carried out under aseptic conditions as described previously [19,35]. Each marmoset was placed in a stereotaxic instrument (SR-6C-HT; Narishige, Tokyo, Japan), and anesthesia was maintained with 2.0–2.5% isoflurane at a flow rate of 1 L/min. Percutaneous oxygen saturation (SpO_2_), pulse rate, and rectal temperature were monitored throughout surgery. In the perioperative period, ampicillin (16.7 mg/kg; antibiotic), carprofen (4.4 mg/kg; anti-inflammatory agent), dexamethasone (1 mg/kg; corticosteroid), atropine (0.1 mg/kg; anticholinergic agent), and maropitant (1 mg/kg; antiemetic) were administered intramuscularly. D-mannitol (10 mL/kg), acetated Ringer’s solution (10 mL/kg), and riboflavin sodium phosphate (200 mg/kg) were administered subcutaneously.

For implantation of the head-fixation plate, the scalp was depilated, sterilized, and incised after topical application of lidocaine, and the connective tissue was removed to expose the skull. After six to seven small screws were anchored to the skull, the head-fixation plate was attached to the skull with universal primer (Tokuyama Dental, Tokyo, Japan), dual-cured adhesive resin cement (Bistite II or Estecem II; Tokuyama Dental), and dental resin cement (Super-Bond; Sun Medical, Shiga, Japan).

For probe implantation, a craniotomy was made over the insertion site, and the dura was incised. A Neuropixels 1.0 probe (PRB_1_4_0480_1_C; IMEC, Leuven, Belgium) [36] was inserted into the brain using a manipulator. A silver wire was placed between the skull and the dura as a reference electrode and connected to the reference and ground wires of the probe. The probe was secured to the head-fixation plate with dental resin via a custom 3D-printed sleeve, and the area around the probe and the entire skull surface were protected by a 3D-printed case and conductive tape for electrostatic shielding.

### Postoperative CT and localization of probe trajectories

At least 1 week after probe implantation, postoperative CT images were acquired (see CT and MR image acquisition). The probe trajectory was approximated by a straight line on the postoperative CT, and points were placed along this line at 10-µm intervals from the probe tip. Each point was mapped into atlas space using the consensus mapping obtained by rigid registration with the Lower set (*N*_ref_ = 6) and was assigned the atlas label of the nearest voxel. The anatomical location of each recording site was estimated on the basis of its distance from the probe tip.

### LFP recording and analysis

Recordings were performed in awake, head-free marmosets whose bodies were restrained with a jacket using a previously described system [19]. Signals were acquired using the Open Ephys GUI (v1.0.1; https://open-ephys.org) [37] with the Neuropix-PXI plugin (v1.0.3). Local field potential (LFP)-band signals sampled at 2,500 Hz were used for analysis and were re-referenced by subtracting the median across all 384 channels from each channel at each time point.

To exclude channels contaminated by external noise, Pearson’s correlation coefficient between each channel and a recording site located outside the skull was computed using LFP signals downsampled to 250 Hz, and 73 channels with *r* > 0.7 were excluded. Sampling depths were set at 60-µm intervals from 0 to 6,865 µm above the probe tip. At each sampling depth, the remaining channel within ±30 µm that was closest to a lateral position of 32 µm was selected as the representative channel, and depths without such channels were treated as missing (black bands).

Correlations between channels in the 8–30 Hz band were estimated as band-limited zero-lag correlations in the frequency domain. The entire re-referenced recording was divided into non-overlapping 10-s segments. In each segment, the mean of each channel was subtracted, a Hann window was applied, and the signals were Fourier transformed. For frequency components with 8 ≤ *f* < 30 Hz, the real part of the cross-spectrum between each pair of channels and the power of each channel were accumulated across all segments. The correlation coefficient was obtained by normalizing the accumulated cross-spectrum by the square root of the product of the accumulated powers of the two channels.

### Statistical analysis

Image processing was performed using 3D Slicer 5.8.1 (https://www.slicer.org) [38] and Python (Python Software Foundation, DE, USA), and image registration was performed using ANTs 2.5.3.post10 (https://github.com/ANTsX/ANTs). LFP and statistical analyses were performed using MATLAB R2025a (MathWorks, MA, USA). The target marmoset was used as the statistical unit, and data are presented as mean ± standard deviation (SD) unless otherwise specified. Within each transformation model and error measure, each landmark set was compared with intensity-based registration using paired *t*-tests with Bonferroni correction for three comparisons (Figures 3 and 4B– E). Errors with *N*_ref_ = 1 and 5 were compared using paired *t*-tests (Figure 4A), and the importance of each landmark was compared with zero using one-sample *t*-tests (Figure 5B). Mapping errors obtained with the raters’ landmarks were compared with those obtained with the original landmarks using paired *t*-tests with Bonferroni correction for the two raters (Figure 5D). The null hypothesis was rejected at the two-sided *P* < 0.05 level. Results of all statistical tests are summarized in Supplementary Table S1. Box plots show the median, interquartile range, and range. In the spatial profiles, lines and shaded areas represent the mean and the mean ± 2 standard errors of the mean (SEM), respectively.

**Figure 3.**
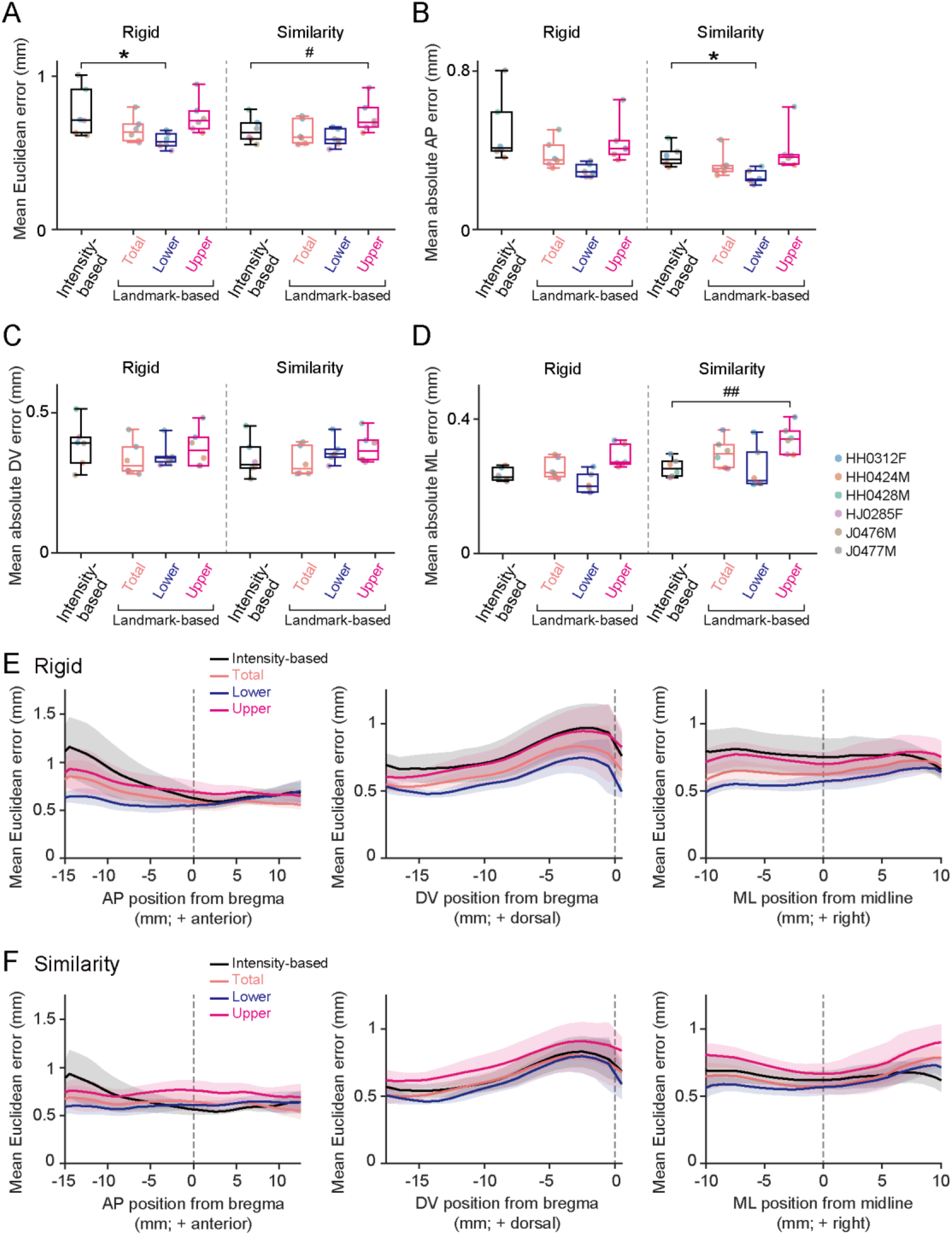
Comparison of CT-to-atlas mapping accuracy across registration methods using a single reference marmoset. (A) Mean Euclidean mapping errors of intensity-based CT-to-CT registration and landmark-based registration with the Total, Lower, and Upper landmark sets under the rigid (left) and similarity (right) transformation models (*n* = 6 marmosets). Rigid, Lower versus intensity-based, *t*(5) = −3.70, \**P* = 0.042; similarity, Upper versus intensity-based, *t*(5) = 4.16, #*P* = 0.026, paired *t*-test with Bonferroni correction. (B–D) Same as A but for mean absolute errors along the anteroposterior (AP; B), dorsoventral (DV; C), and mediolateral (ML; D) axes. B, similarity, Lower versus intensity-based, *t*(5) = −5.23, \**P* = 0.010; D, similarity, Upper versus intensity-based, *t*(5) = 6.68, ^#*#*^*P* = 0.0034. In A–D, each dot represents a target marmoset, with colors indicating individual animals, and * and # indicate errors significantly smaller and larger, respectively, than those of intensity-based registration (*\*P* or *^#^P* < 0.05; *^##^P* < 0.01). (E, F) Mean Euclidean mapping errors plotted against AP (left), DV (middle), and ML (right) positions for the rigid (E) and similarity (F) transformation models. Vertical dashed lines indicate zero along each axis.

**Figure 4.**
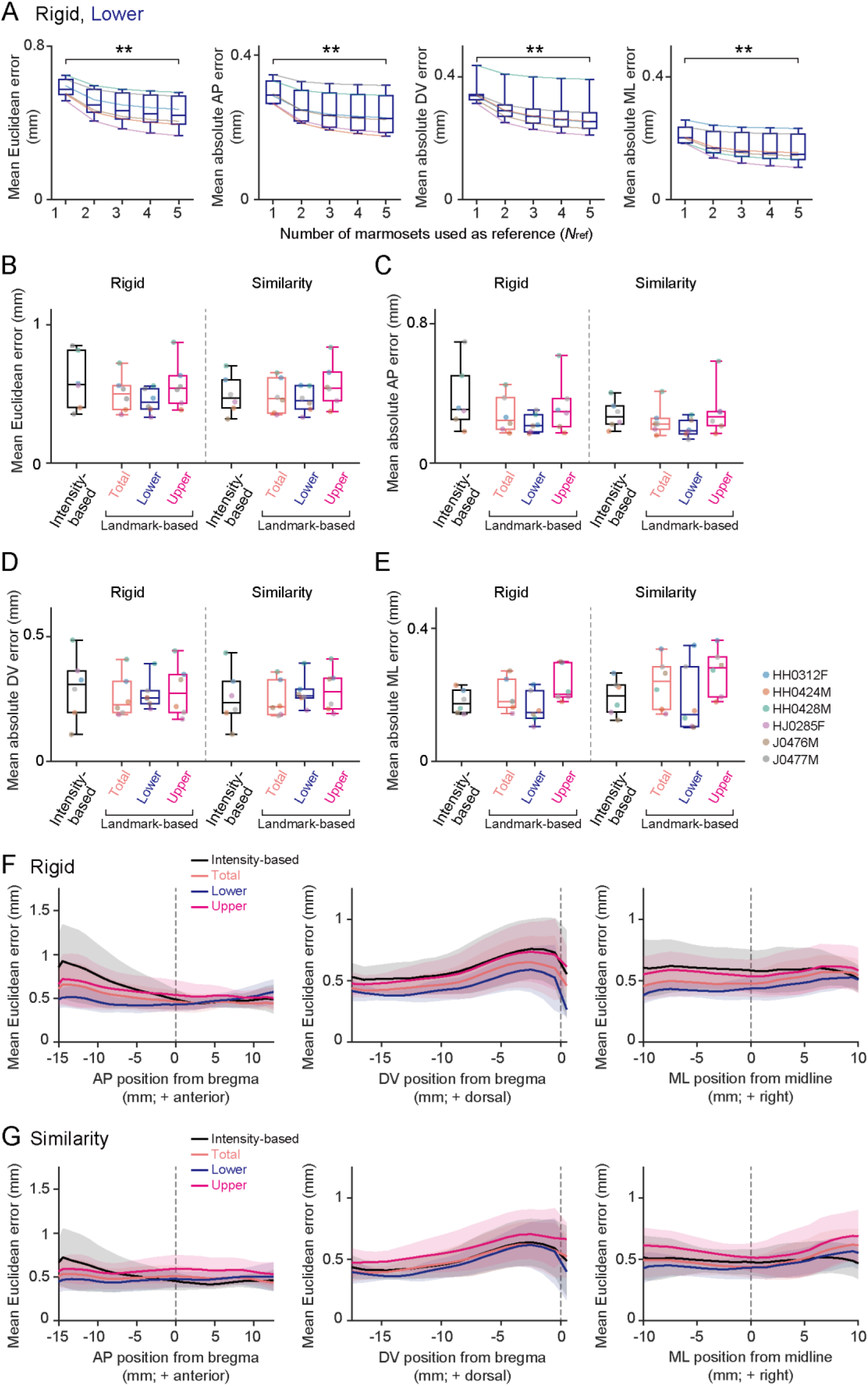
Using multiple reference marmosets improves CT-to-atlas mapping accuracy. (A) Mean Euclidean mapping errors and mean absolute errors along the anteroposterior (AP), dorsoventral (DV), and mediolateral (ML) axes (from left to right) as a function of the number of reference marmosets (*N*_ref_ = 1–5) for rigid landmark-based registration with the Lower landmark set (*n* = 6 marmosets). *N*_ref_ = 1 versus *N*_ref_ = 5: Euclidean, *t*(5) = 8.39, \*\**P* = 0.00039; AP, *t*(5) = 6.51, \*\**P* = 0.0013; DV, *t*(5) = 8.81, \*\**P* = 0.00031; ML, *t*(5) = 5.79, \*\**P* = 0.0022, paired *t*-test. (B–E) Mean Euclidean mapping errors (B) and mean absolute errors along the AP (C), DV (D), and ML (E) axes of intensity-based CT-to-CT registration and landmark-based registration with the Total, Lower, and Upper landmark sets with five reference marmosets (*N*_ref_ = 5) under the rigid (left) and similarity (right) transformation models (*n* = 6 marmosets). Each line (A) or dot (B–E) represents a target marmoset, with colors indicating individual animals. (F, G) Mean Euclidean mapping errors with five reference marmosets plotted against AP (left), DV (middle), and ML (right) positions for the rigid (F) and similarity (G) transformation models. Vertical dashed lines indicate zero along each axis.

**Figure 5.**
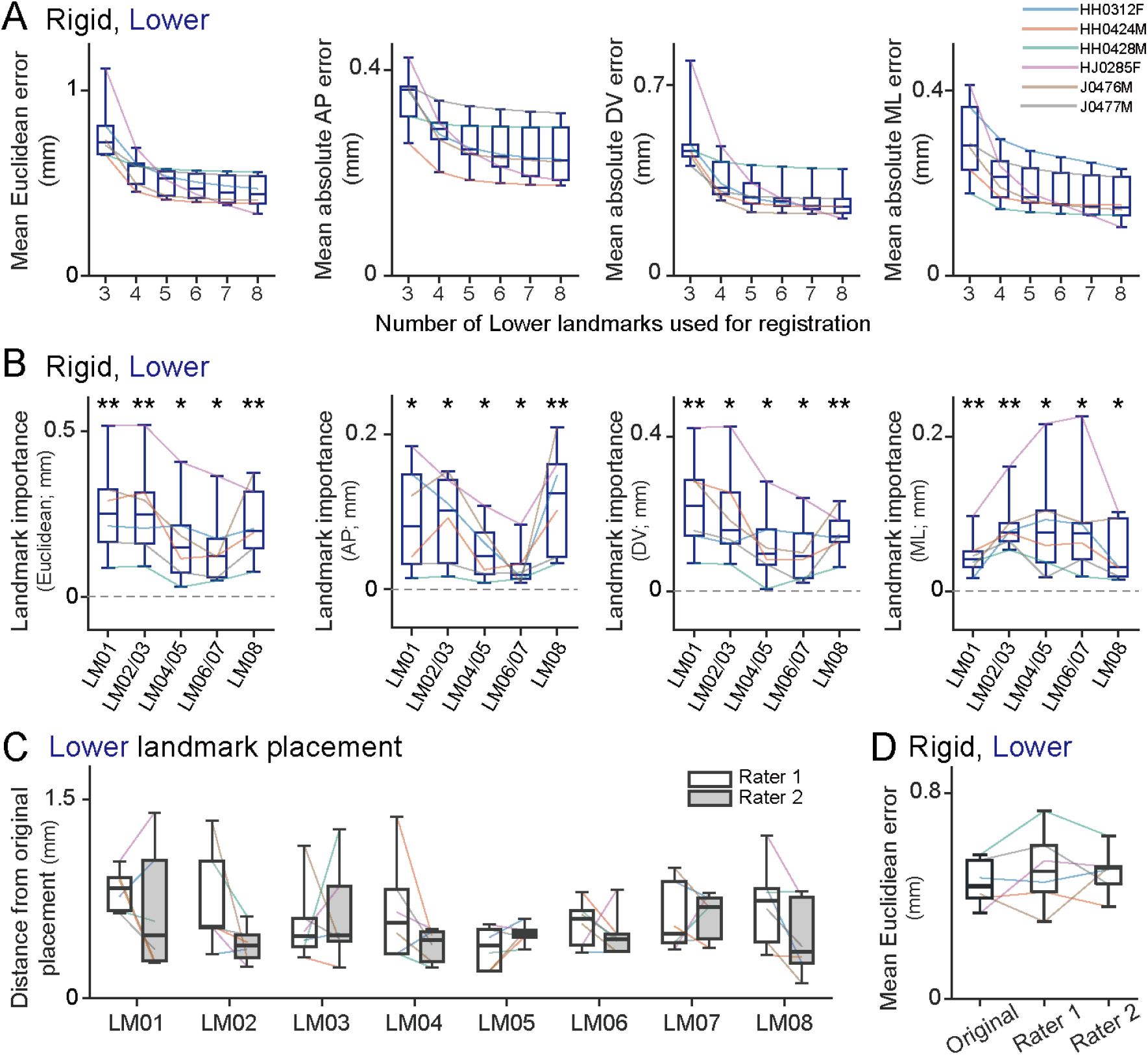
Sensitivity of CT-to-atlas mapping accuracy to the number, identity, and placement of cranial base landmarks. All mapping errors were obtained using rigid registration with the Lower landmark set and five reference marmosets (*N*_ref_ = 5). (A) Mean Euclidean mapping errors and mean absolute errors along the anteroposterior (AP), dorsoventral (DV), and mediolateral (ML) axes (from left to right) as a function of the number of landmarks used for registration (*n* = 6 marmosets). For each number of landmarks, errors were averaged across all combinations selected from the eight Lower landmarks. (B) Landmark importance, defined as the reduction in mapping error upon adding the indicated landmark to three-landmark base configurations not containing it, for the Euclidean, AP, DV, and ML errors (from left to right; *n* = 6 marmosets). Values for bilateral landmarks (LM02/03, LM04/05, and LM06/07) were averaged across the left and right counterparts. Dashed lines indicate zero. Asterisks indicate importance values significantly different from zero (one-sample *t*-test; \**P* < 0.05, \*\**P* < 0.01). (C) Distances between the Lower landmarks placed by two raters and the original landmarks. (D) Mean Euclidean mapping errors with the original landmarks and with those placed by each rater. In A–D, each line represents a target marmoset, with colors indicating individual animals.

## Results

### Construction of a registration pathway from CT images to the brain atlas

To map CT images to the brain atlas without requiring MRI of the target marmoset, we constructed a registration pathway via a reference marmoset for which both CT and MR images were available (Figure 1A). The pathway consisted of three steps: skull CT registration from the target to the reference marmoset, rigid CT-to-MR registration within the reference marmoset, and registration of the reference MR image to the T2-weighted atlas template [24]. The transformations obtained from these steps were concatenated to map the CT coordinates of the target marmoset into the BMA 2.0 space.

For interindividual skull CT registration, we compared two approaches [39]: intensity-based registration, which aligns CT images according to their signal intensity, and landmark-based registration, which aligns them according to corresponding anatomical landmarks. For landmark-based registration, we defined 13 skull landmarks (LM01–13) and grouped them into three landmark sets: Total, comprising all 13 landmarks; Lower, comprising eight landmarks located mainly on the cranial base (LM01–08); and Upper, comprising eight landmarks including those on the calvaria (LM06–13) (Figure 1B).

### CT-to-atlas mapping accuracy using a single reference marmoset

To quantify mapping accuracy, we used six marmosets for which both CT and MR images were available. For each target marmoset, a grid of points defined within the intracranial area of the CT image was mapped into atlas space through two pathways: within-subject mapping via the target’s own MR image (Figure 2, upper), and cross-subject mapping via the CT and MR images of a reference marmoset (Figure 2, lower). The within-subject mapping served as the benchmark for evaluating the cross-subject mapping, which represents the proposed pathway. Mapping error was defined as the Euclidean distance between the atlas-space coordinates of corresponding grid points obtained through the two pathways and was averaged across grid points. The MR image of the target marmoset was used only for the within-subject mapping and not for the cross-subject mapping.

We first evaluated mapping accuracy with a single reference marmoset (*N*_ref_ = 1). For each target marmoset, mapping errors were computed using each of the remaining five marmosets as the reference and then averaged across the five references (Figure 3). We compared intensity-based registration with landmark-based registration using the Total, Lower, and Upper landmark sets under the rigid and similarity transformation models. With the rigid model, the Lower set yielded a significantly smaller mean Euclidean error than intensity-based registration (Figure 3A, left; Lower, 0.58 ± 0.05 mm; intensity-based, 0.77 ± 0.16 mm; *n* = 6, *t*(5) = −3.70, *P* = 0.042, paired *t*-test with Bonferroni correction), whereas the Total and Upper sets did not (Figure 3A, left; Total, 0.65 ± 0.09 mm, *n* = 6, *t*(5) = −2.18, *P* = 0.24; Upper, 0.74 ± 0.11 mm, *n* = 6, *t*(5) = −0.44, *P* > 0.99). With the similarity model, the mean Euclidean errors of the Total and Lower sets did not differ significantly from that of intensity-based registration (Figure 3A, right; intensity-based, 0.65 ± 0.08 mm, *n* = 6; Total, 0.63 ± 0.08 mm, *n* = 6, *t*(5) = −1.02, *P* > 0.99; Lower, 0.60 ± 0.06 mm, *n* = 6, *t*(5) = −2.36, *P* = 0.19), whereas that of the Upper set was significantly larger (0.74 ± 0.11 mm, *n* = 6, *t*(5) = 4.16, *P* = 0.026).

Decomposition of the errors into the anteroposterior (AP), dorsoventral (DV), and mediolateral (ML) components showed that, with the similarity model, the Lower set had a significantly smaller mean absolute AP error (Figure 3B; intensity-based, 0.37 ± 0.05 mm; Lower, 0.27 ± 0.03 mm; *n* = 6, *t*(5) = −5.23, *P* = 0.010, paired *t*-test with Bonferroni correction) and the Upper set had a significantly larger mean absolute ML error (Figure 3D; intensity-based, 0.26 ± 0.03 mm; Upper, 0.34 ± 0.04 mm; *n* = 6, *t*(5) = 6.68, *P* = 0.0034) than intensity-based registration. No significant differences in mean absolute DV error were found between landmark-based and intensity-based registration with either model (Figure 3C; *n* = 6, *t*(5) ranging from −2.03 to 1.91, *P* > 0.29 for all comparisons).

We next examined how mapping error varied with position in the brain by plotting the mean Euclidean error along the AP, DV, and ML axes. With the rigid model, intensity-based registration and the Total and Upper sets showed increased errors in posterior regions, whereas the Lower set showed a smaller increase and a relatively uniform error profile along the AP axis (Figure 3E). With the similarity model, the posterior increase in error observed with intensity-based registration was likewise smaller for the Lower set (Figure 3F). Although the error of the Lower set, like that of intensity-based registration, increased toward dorsal positions, it remained generally low across the AP, DV, and ML profiles (Figure 3E, F).

### Reduction of mapping error by using multiple reference marmosets

We then examined whether combining the mappings obtained via multiple reference marmosets reduces mapping error (Figure 4). With each of the six marmosets used in turn as the target, the number of reference marmosets selected from the remaining five was increased from one to five (*N*_ref_ = 1–5). For each point in the target CT, the atlas-space coordinates obtained via the individual references were averaged with equal weights, and mapping errors were calculated for all possible combinations of reference marmosets and then averaged within each target marmoset. With the rigid model and the Lower set, the group means of the mean Euclidean error and the mean absolute AP, DV, and ML errors decreased monotonically with increasing *N*_ref_, and all errors were significantly smaller with *N*_ref_ = 5 than with *N*_ref_ = 1 (Figure 4A; mean Euclidean error: *N*_ref_ = 1, 0.58 ± 0.05 mm; *N*_ref_ = 5, 0.45 ± 0.09 mm; *n* = 6, *t*(5) = 8.39, *P* = 0.00039; mean absolute AP, DV, and ML errors: *t*(5) = 6.51, 8.81, and 5.79, *P* = 0.0013, 0.00031, and 0.0022, respectively, paired *t*-test). These reductions corresponded to 22.5%, 20.8%, 22.6%, and 22.8% of the errors with *N*_ref_ = 1, respectively. Under the other registration conditions, the mean Euclidean error likewise decreased monotonically with increasing *N*_ref_, with reductions of 22.6–23.9% from *N*_ref_ = 1 to 5.

With five reference marmosets (*N*_ref_ = 5), landmark-based and intensity-based registration did not differ significantly in mean Euclidean error or in mean absolute AP, DV, or ML error with either the rigid or similarity model (Figure 4B–E; *n* = 6, *t*(5) ranging from −2.59 to 2.65, *P* > 0.13 for all comparisons, paired *t*-test with Bonferroni correction). Nevertheless, the Lower set showed a smaller group-mean Euclidean error than intensity-based registration with both models (Figure 4B; rigid: intensity-based, 0.59 ± 0.21 mm; Lower, 0.45 ± 0.09 mm; *n* = 6, *t*(5) = −2.41, *P* = 0.18; similarity: intensity-based, 0.49 ± 0.14 mm; Lower, 0.46 ± 0.09 mm; *n* = 6, *t*(5) = −0.96, *P* > 0.99), and a similar tendency was observed for the mean absolute AP error (Figure 4C; rigid: intensity-based, 0.39 ± 0.20 mm; Lower, 0.24 ± 0.06 mm; *n* = 6, *t*(5) = −2.56, *P* = 0.15; similarity: intensity-based, 0.29 ± 0.08 mm; Lower, 0.21 ± 0.05 mm; *n* = 6, *t*(5) = −2.59, *P* = 0.15).

The position-dependent error profiles showed the same pattern as with a single reference marmoset: intensity-based registration showed increased errors in posterior regions, whereas this increase was smaller for the Lower set (Figure 4F, G). The Lower set also showed generally low errors along the DV and ML axes, and with the rigid model in particular, its error tended to be lower than that of intensity-based registration over a wide range of positions (Figure 4F). Thus, the use of multiple reference marmosets reduced mapping error, and the Lower set maintained relatively low errors under this condition, as it did with a single reference marmoset.

### Effects of landmark number and contributions of individual cranial base landmarks

Using rigid registration with the Lower set and five reference marmosets (*N*_ref_ = 5), we next varied the number of landmarks used for registration from three to eight. Averaged across all landmark combinations for each number of landmarks, the mean Euclidean error and the mean absolute AP, DV, and ML errors decreased as the number of landmarks increased (mean Euclidean error: three landmarks, 0.78 ± 0.18 mm; eight landmarks, 0.45 ± 0.09 mm; *n* = 6). The largest reduction occurred from three to four landmarks (four landmarks, 0.57 ± 0.08 mm), and further additions yielded smaller reductions; with five or more landmarks, the group-mean Euclidean error differed from that with all eight landmarks by 0.06 mm or less (95% CI of the difference with five landmarks, −0.01 to 0.12 mm) (Figure 5A).

To evaluate the contribution of each landmark, we defined landmark importance as the reduction in error when a landmark was added to three-landmark base configurations that did not include it. For bilateral landmarks (LM02/03, LM04/05, and LM06/07), landmark importance was averaged across the left and right counterparts. Landmark importance was significantly greater than zero for all landmarks and bilateral landmark pairs in all error metrics (Figure 5B; *n* = 6, *t*(5) ranging from 2.58 to 6.34, *P* < 0.05 for all comparisons, one-sample *t*-test) and, for the Euclidean error, was relatively large for LM01 and LM02/03 (LM01, 0.27 ± 0.15 mm; LM02/03, 0.26 ± 0.15 mm). Along the individual axes, LM08 contributed relatively strongly to reducing the AP error (0.12 ± 0.07 mm), and LM01 and LM02/03 to reducing the DV error (0.23 ± 0.13 and 0.20 ± 0.13 mm, respectively). For the ML error, the bilateral landmarks (LM02/03, LM04/05, and LM06/07; 0.09 ± 0.04, 0.09 ± 0.07, and 0.09 ± 0.07 mm, respectively) showed larger importance than the midline landmarks (LM01 and LM08; 0.05 ± 0.03 and 0.05 ± 0.04 mm, respectively), with the most consistent contribution from LM02/03 (Figure 5B).

We next examined whether mapping accuracy depended on the operator. The landmarks placed by two inexperienced raters differed from the original landmarks by 0.38–0.82 mm on average (Figure 5C). Nevertheless, the mean Euclidean error with rigid registration increased by only approximately 0.05 mm (original, 0.45 ± 0.09 mm; rater 1, 0.51 ± 0.15 mm; rater 2, 0.49 ± 0.09 mm; *P* > 0.5; Figure 5D), with similar results for similarity registration (Supplementary Table S1).

### Application to CT data acquired with a different scanner

To assess the applicability of the registration pathway to CT data acquired under different imaging conditions, we mapped three marmosets scanned at another facility (UT1, UT2, and UT3; Table 1), for which both CT and MR images were available, using all six marmosets of the evaluation dataset as references (*N*_ref_ = 6). Although these CT images differed substantially in resolution from those of the evaluation dataset, the corresponding cranial base landmarks could be identified (Figure 6A). With the rigid model, the Lower set yielded smaller mean Euclidean errors and mean absolute AP, DV, and ML errors than intensity-based registration in all three target marmosets (Figure 6B, C; mean Euclidean error: intensity-based, 0.71–1.36 mm; Lower, 0.61–0.69 mm; *n* = 3). Thus, multi-reference mapping with cranial base landmarks was applicable to CT data acquired with a different scanner and resolution, and the Lower set again yielded lower errors than intensity-based registration.

**Figure 6.**
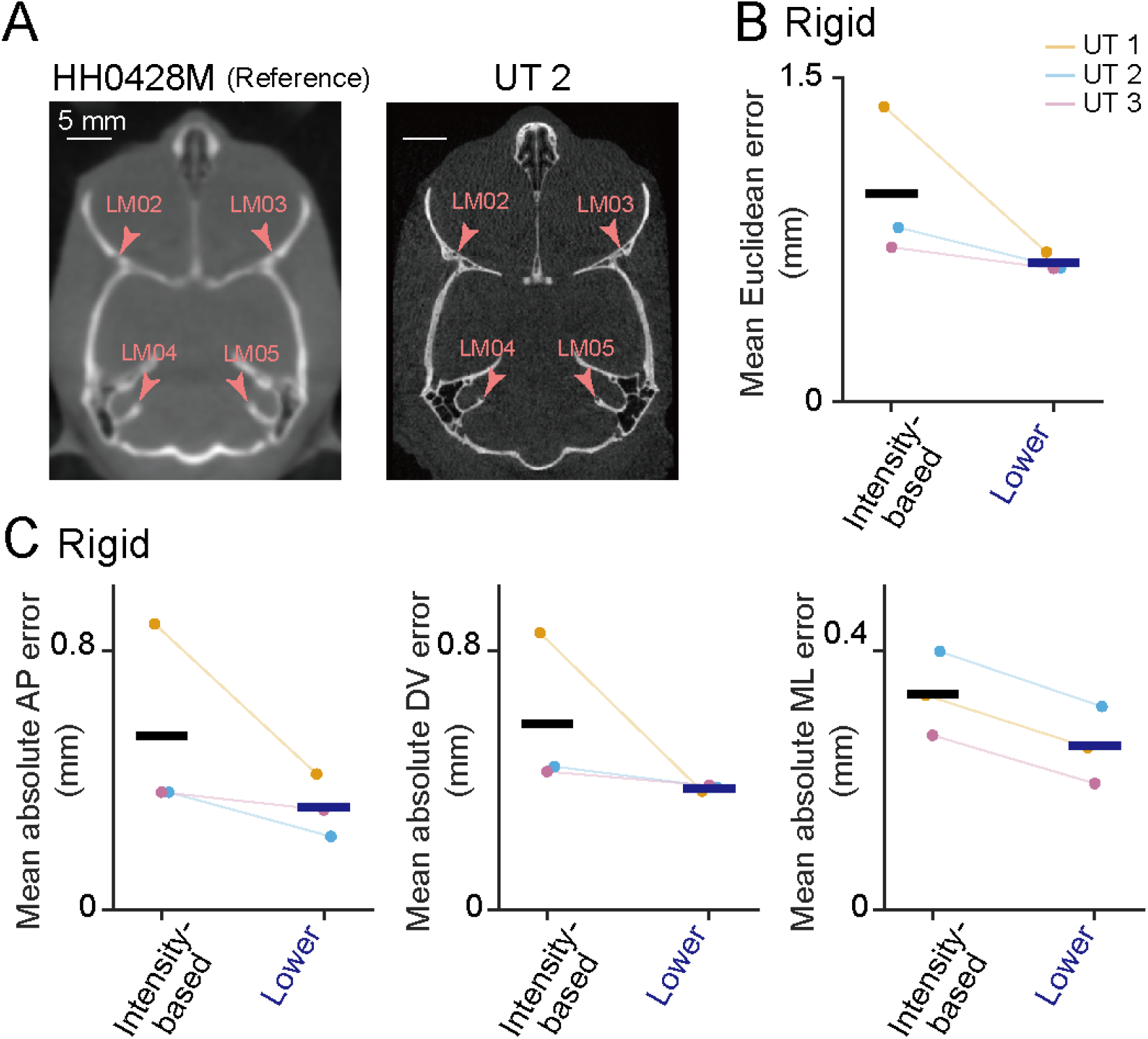
Application of registration based on cranial base landmarks to CT data acquired with a different scanner. (A) Representative horizontal CT sections from one of the reference marmosets (HH0428M; left; voxel size, 0.25 × 0.25 × 0.5 mm) and a target marmoset scanned with a different CT scanner at a higher resolution (UT2; right; voxel size, 0.037 × 0.037 × 0.037 mm). Pink arrowheads indicate four corresponding landmarks of the Lower set (LM02–05). Scale bars, 5 mm. (B, C) Mean Euclidean mapping errors (B) and mean absolute errors along the anteroposterior (AP), dorsoventral (DV), and mediolateral (ML) axes (C, from left to right) of intensity-based registration and landmark-based registration with the Lower landmark set (*n* = 3 marmosets). Rigid registration was used with all six marmosets of the evaluation dataset as references (*N*_ref_ = 6). In B and C, each line represents a target marmoset, with colors indicating individual animals, and horizontal bars indicate means.

### Marmoset Atlas Mapper: open-source software for multi-reference CT-to-atlas mapping

We implemented the multi-reference registration pathway in Marmoset Atlas Mapper, a new open-source software package (Figure 7). The user loads a target CT image and selects the registration type and transformation model (Figure 7A). Landmarks are then placed on the target CT, with the corresponding landmarks on a reference CT displayed as a guide (Figure 7B). Because landmarks are matched to their counterparts in the reference CTs by label, registration can also be performed when only a subset of the landmarks is placed. When the registration is run, the target CT is registered to the CT of each of the six reference marmosets in the evaluation dataset, and the resulting mappings are averaged (*N*_ref_ = 6). The alignment between the target CT and the atlas MR image can then be inspected in the quality-check window (Figure 7C). The target-to-atlas transformation and its inverse, the atlas images and region labels resampled into target CT space, the target CT resampled into atlas space, and the quality-check results are saved automatically.

**Figure 7.**
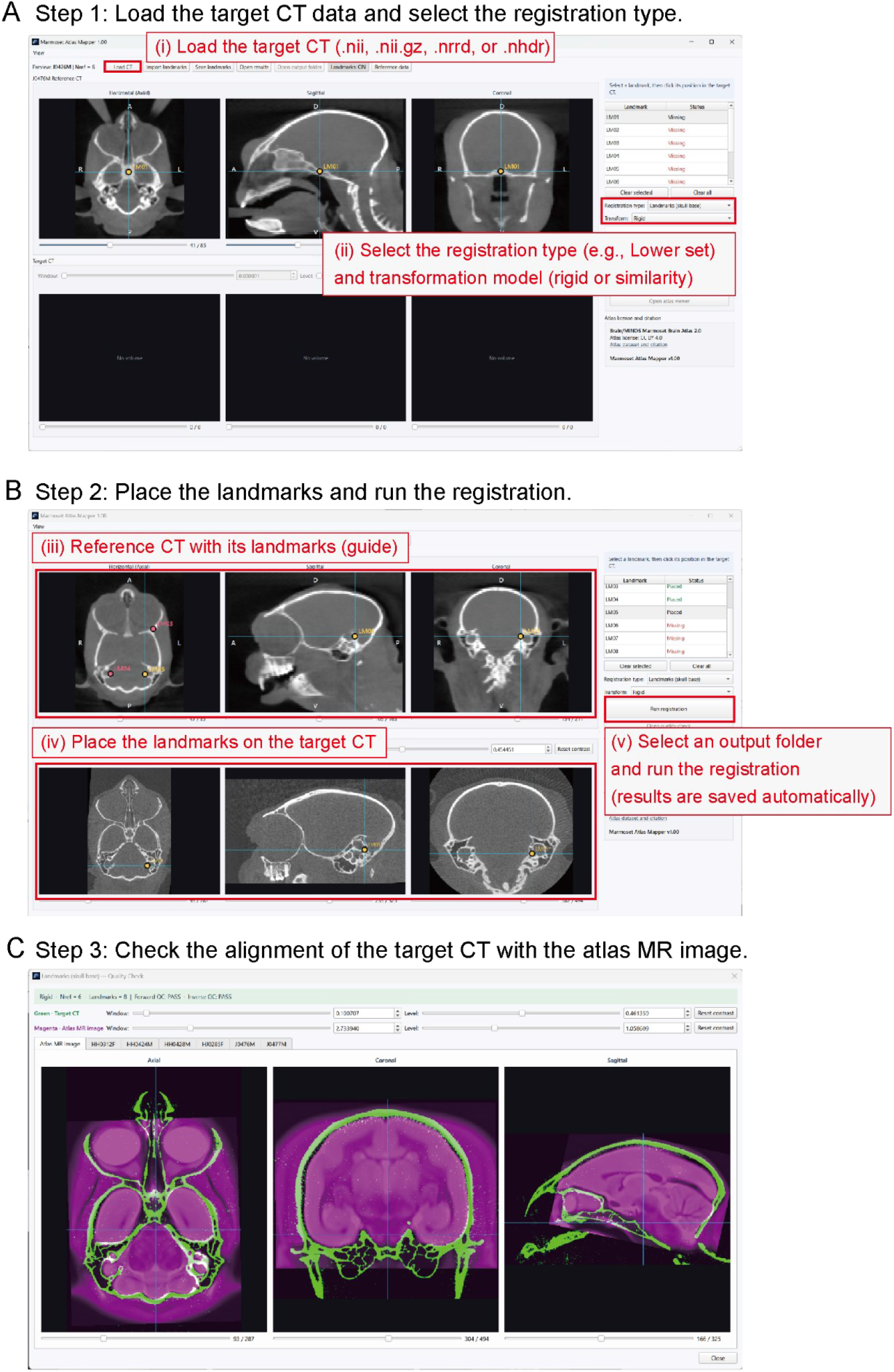
Marmoset Atlas Mapper, an open-source software package for multi-reference CT-to-atlas mapping. Graphical user interface of Marmoset Atlas Mapper (version 1.00). (A) Main window before the target CT is loaded, showing a reference CT (J0476M; top) and an empty target CT view (bottom). The target CT is loaded (i), and the registration type and transformation model are selected (ii). (B) Main window after the target CT (UT1) is loaded. Landmarks are placed on the target CT (iv, bottom), guided by the corresponding landmarks on the reference CT (iii, top), and the registration is run after an output folder is selected (v). (C) Quality-check window showing the target CT (green) overlaid with the atlas MR image (magenta) after registration (Lower set, rigid model, *N*_ref_ = 6).

### Application to the localization of Neuropixels recording sites

We then used Marmoset Atlas Mapper to estimate the locations of Neuropixels implanted in two marmosets (UT4 and UT5; Table 1; Figure 8A). The postoperative CT images contained artifacts from the head-fixation plate, dental cement, and the probe. With rigid registration using all six marmosets of the evaluation dataset as references (*N*_ref_ = 6), intensity-based registration produced visible gaps and overlaps between the atlas brain boundary and the skull contour on CT (Figure 8B, top). Such misalignment was observed in all three postoperative CT scans obtained from two Neuropixels-implanted marmosets. In contrast, with landmark-based registration using the Lower set, the atlas brain boundary matched the skull contour on visual inspection (Figure 8B, bottom).

**Figure 8.**
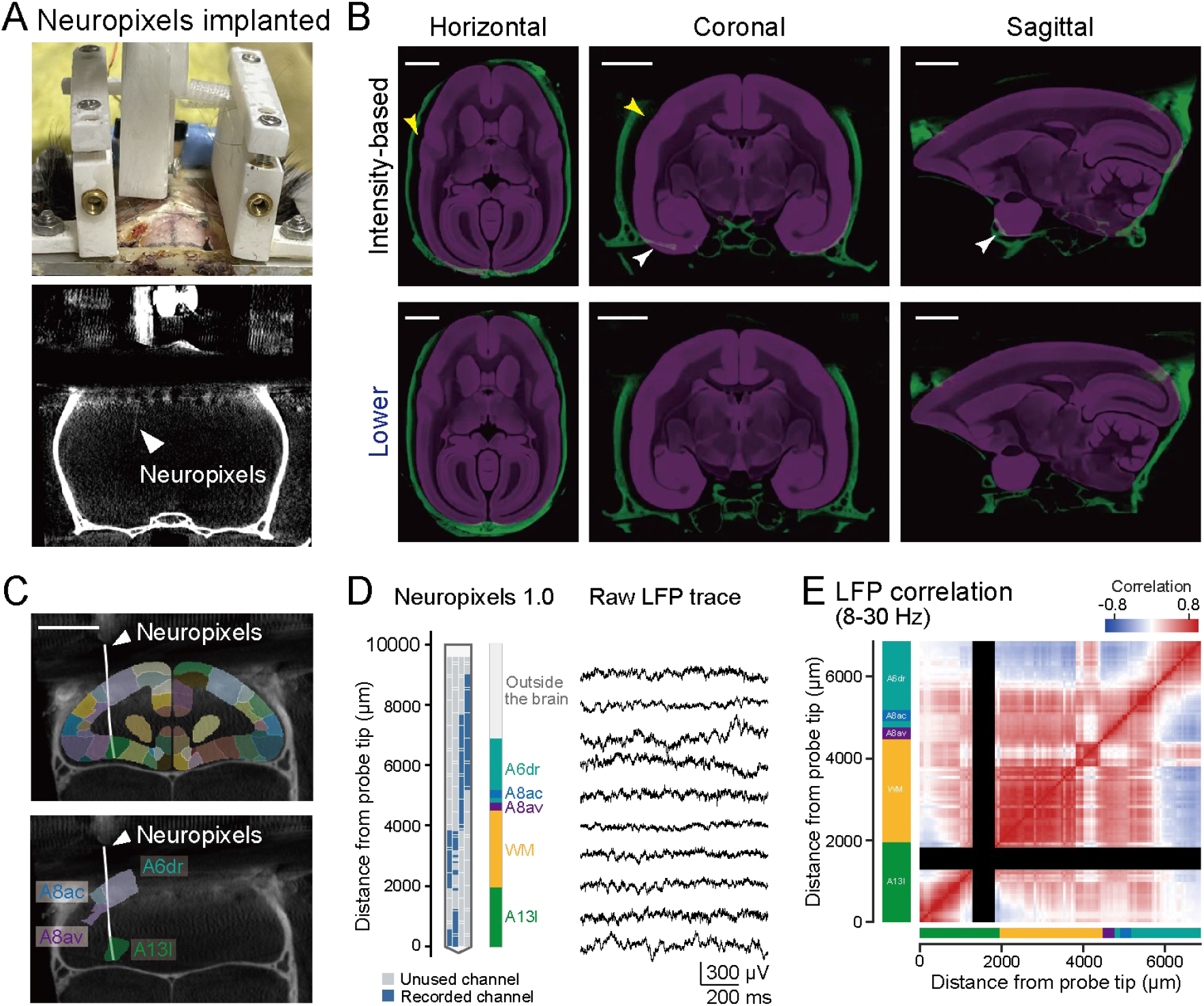
Application of Marmoset Atlas Mapper to postoperative CT for localizing Neuropixels recording sites. (A) Head-fixation plate and an implanted Neuropixels probe (top), and a coronal section of the postoperative CT (bottom). The white arrowhead indicates the probe shank. (B) Horizontal (left), coronal (middle), and sagittal (right) sections of the postoperative CT (green) overlaid with the atlas MR image (magenta) after intensity-based registration (top) and landmark-based registration with the Lower landmark set (bottom). Both registrations used the rigid model with six reference marmosets (*N*_ref_ = 6). Yellow and white arrowheads indicate gaps and overlaps, respectively, between the atlas brain boundary and the skull contour on CT after intensity-based registration. Scale bars, 5 mm. (C) Coronal sections of the postoperative CT in atlas space, overlaid with atlas brain regions (top) and with the regions traversed by the probe (bottom) after landmark-based registration with the Lower landmark set. White lines indicate the probe trajectory identified on CT. Scale bar, 5 mm. (D) Electrode sites of the Neuropixels 1.0 probe (left), their anatomical assignments based on the mapping in C (middle), and representative LFP traces at selected depths (right). (E) Correlation matrix of 8–30 Hz LFP signals between recording channels, ordered by distance from the probe tip. Colored bars along the axes indicate the anatomical assignments in D. Black bands indicate depths with no valid recording channels. WM, white matter; other abbreviations denote brain regions defined in BMA 2.0.

Using the mapping obtained with the Lower set, we assigned atlas-defined brain regions along the probe trajectory identified on CT (Figure 8C). According to this mapping, the probe passed, from the brain surface, through A6dr, A8ac, A8av, the white matter (WM), and A13l (Figure 8D). We further computed the correlations of 8–30 Hz local field potential (LFP) signals between recording channels and found that the transitions in the correlation structure were qualitatively consistent with the atlas-defined boundaries between the dorsal cortex (A6dr–A8av), WM, and the ventral cortex (A13l) (Figure 8E). These results illustrate how atlas mapping based on cranial base landmarks with Marmoset Atlas Mapper can be used to estimate the brain regions of individual recording sites from postoperative CT.

## Discussion

In this study, we constructed a registration pathway that maps marmoset CT images into the BMA 2.0 space without requiring MRI of the target marmoset. In this pathway, the CT image of the target marmoset is registered to that of a reference marmoset for which both CT and MR images are available, and the resulting transformation is concatenated with the CT-to-MR and MR-to-atlas transformations of the reference marmoset. With a single reference marmoset, rigid registration using cranial base landmarks (the Lower set) yielded a significantly smaller mapping error than intensity-based registration. Averaging the mappings obtained via five reference marmosets reduced the error by approximately 22–24% relative to a single reference under all registration conditions, yielding a mean Euclidean error of 0.45 mm with the Lower set and the rigid model. The pathway could also be applied to CT images acquired with a different scanner and to postoperative CT images of implanted marmosets. This pathway is implemented in Marmoset Atlas Mapper, our newly developed open-source software. As an application, we used it to estimate the atlas-defined brain regions of Neuropixels recording sites. To our knowledge, this is the first study in marmosets to quantify the accuracy of CT-to-atlas mapping against the within-subject mapping through each animal’s own MR image and to compare registration methods for this purpose.

Marmosets are outbred and genetically diverse, and, as noted in the Introduction, they show substantial interindividual variability in the shape and size of the skull and brain [14,21]. This variability is particularly pronounced in the calvaria. The anteroposterior (AP) position of bregma relative to the brain varies by approximately 1 mm (SD), three to four times as much as in laboratory rodents even after normalization for brain size, and the contour of the calvaria varies considerably across animals despite alignment of the skulls on the external auditory canals and infraorbital margins [14]. The marked dependence of mapping error on the choice of landmarks in the present study likely reflects these regional differences in the magnitude of interindividual variability across the skull. Consistent with this interpretation, the Upper set, which includes landmarks on the calvaria, and intensity-based registration, which aligns entire images including this region, yielded larger errors than the Lower set, and their errors increased toward posterior regions. In contrast, the Lower set showed a relatively uniform error profile along the AP axis. Unlike the posterior increase, an increase in error toward dorsal positions occurred with both the Lower set and intensity-based registration. Because this dorsal increase was common to both registration approaches, it is likely attributable, at least in part, to interindividual differences in dorsal brain shape that cannot be accommodated by either rigid or similarity transformations. Adding isotropic scaling (the similarity model) did not reduce the error of the Lower set; we therefore adopted the simpler rigid model.

In the evaluation dataset, in which all CT images were acquired with the same scanner, mapping errors with five reference marmosets (*N*_ref_ = 5) did not differ significantly between the Lower set and intensity-based registration. However, in the external dataset, whose CT images were acquired with a different scanner and at a different resolution, the Lower set yielded smaller errors than intensity-based registration in all three marmosets. Moreover, on postoperative CT images, intensity-based registration produced visible misalignment between the atlas brain boundary and the skull contour. Because the intensity-based registration used in this study depends on intensity similarity across entire images [39], it is susceptible to differences in resolution and field of view between images, as well as to artifacts caused by head-fixation plates, dental cement, and electrodes [40]. In contrast, landmark-based registration relies solely on corresponding anatomical points on the skull and is therefore less affected by these factors, provided that the landmarks can be identified. Furthermore, because head-fixation plates and electrodes are typically mounted on the calvaria, cranial base landmarks remain identifiable on postoperative CT images. These features make registration based on cranial base landmarks particularly well suited to postoperative images and to images acquired with different scanners or settings. In addition, when five or more of the eight Lower landmarks were used, the error averaged across all combinations was within 0.06 mm of that obtained with all eight, suggesting that mapping accuracy is largely maintained even when some landmarks cannot be identified, for example because of image artifacts or a limited field of view. However, the bilateral landmarks (LM02/03, LM04/05, and LM06/07) contributed more than the midline landmarks (LM01 and LM08) to reducing the mediolateral (ML) error. This is likely because the midline landmarks alone poorly constrain rotation about the AP axis (i.e., left–right tilt). Therefore, when the number of landmarks is reduced, it is important to retain landmarks that are separated from one another in different directions. Although the landmarks placed by two inexperienced raters differed from the original landmarks by 0.4–0.8 mm, the mapping error increased by only approximately 0.05 mm, likely because placement errors of individual landmarks are partly averaged out by fitting to multiple landmarks and by averaging across references. This suggests that users without prior experience can achieve comparable accuracy given the landmark definitions.

High-density probes such as Neuropixels have also been adopted in nonhuman primates [41] and allow simultaneous recording from multiple regions along a single insertion track. Accordingly, all recording sites along the track must be localized in three dimensions, including those in deep structures. In addition, experiments that sample many sites through repeated insertions [7,42,43] benefit from the ability to estimate recording sites *in vivo*. Histological verification is the standard approach [20], but it can be performed only at the end of the experiment. *In vivo*, recording sites can be localized precisely by integrating CT with each animal’s MR images [14], but this requires acquiring MR images of every animal, which entails access to imaging facilities as well as the time and cost of imaging. Among approaches that do not require subject-specific MRI, stereotaxic coordinates, which are standard in rodents, provide limited accuracy in marmosets because of the cranial variability described above [14]. Another approach fits an MRI-based template brain, such as Marmoset Brain Mapping V3 [21], to the skull reconstructed from each animal’s CT to estimate target and recording locations [7,12]; however, it has been used mainly to infer the cortical areas beneath the insertion sites, and its accuracy has not been quantified. In contrast, because the present method maps CT images into three-dimensional atlas space without requiring subject-specific MRI, it allows recording sites to be estimated *in vivo* along the entire insertion track, including deep structures, from the electrode positions visible on CT. Such *in vivo* estimation complements, rather than replaces, postmortem histological verification. This advantage is not limited to electrophysiology. The method could likewise be applied when optogenetic manipulations [9,11–13,44,45] or anterograde and retrograde tracer injections [17,18,46–48] target subcortical as well as cortical structures, and is therefore expected to contribute to the understanding of neural circuits in primates.

This study has two main limitations. First, mapping accuracy was evaluated not against histology but against the within-subject mapping through each target marmoset’s own MR image. Because this within-subject mapping itself contains MR-to-atlas registration error, the mean error of 0.45 mm does not necessarily represent the deviation from the true anatomical location, and assignments of recording sites near regional boundaries or within small structures remain uncertain. Nevertheless, because the same within-subject mapping was used for all registration methods, agreement with this mapping could be compared across methods under identical conditions. In addition, the application to postoperative CT was evaluated by visual inspection, and the correspondence between the assigned regions and the LFP correlation structure was assessed only qualitatively; LFP correlations can also be affected by the referencing scheme and by volume conduction. Second, the evaluation dataset comprised only six marmosets, and the number of reference marmosets was correspondingly limited. In the present study, mapping error decreased monotonically as *N*_ref_ increased from one to five. In multi-atlas segmentation, accuracy has likewise been shown to improve as more atlases are used and when atlases similar to the target are selected [49,50]. Accuracy may therefore be further improved by increasing the number of reference marmosets and selecting, from among them, those whose skull shapes most closely resemble that of the target marmoset. Unlike population-averaged CT and MR templates, which would require nonlinear registration among the reference marmosets, the present approach uses each reference independently, so that references can be added, selected, or weighted for each target without rebuilding a template.

## Acknowledgements

We are grateful to Dr. Masami Kanai and Dr. Naoaki Mizuno (Institute of Science Tokyo), Dr. Hideaki Shimazaki (Kyoto University), and Dr. Yutaka Sakai (Tamagawa University) for their help with the marmoset experiments and for valuable discussions. We also thank Ms. Chie Soai and Ms. Minori Kajiura for placing the cranial base landmarks used to assess the reproducibility of landmark placement. This work was supported by AMED Brain/MINDS 2.0 (JP24wm0625108 to T. E. and M. M.; JP24wm0625212 to K. O.; JP23wm0625001 to M. M. and T. M.; JP24wm0625106 to Y. I.) and by MEXT/JSPS KAKENHI (JP25KJ0107 to H. Y.; JP22K15222 and JP25H01726 to M. K.; JP25H01767 to Y. Hori; JP23H04977 to T. E.; JP22H05160 to M. M.; JP21H05242 and JP23H02589 to Y. I.).

## Author contributions

H. Y. and Y. I. designed the study. H. Y., M. K., Y. N., Y. Hirayama, T. E., K. S., and K. O. acquired and analyzed the CT data. H. Y., Y. Hori, T. E., and K. S. acquired and analyzed the MR data. H. Y. and M. K. acquired and analyzed the electrophysiological data. M. M., T. M., and Y. I. supervised the project. H. Y. and Y. I. prepared the figures and wrote the manuscript, and all authors reviewed the manuscript.

## Conflict of interest

The authors declare no competing interests.

## Data and code availability

The source code of Marmoset Atlas Mapper is available on GitHub (https://github.com/H-Yagishita/Marmoset_CT-Atlas_Mapper) under the MIT License. The complete package of version 1.00 is deposited in Zenodo (https://doi.org/10.5281/zenodo.23683937). The package contains the Windows executable and its source code, the BMA 2.0 atlas [24], and the CT and MR images, landmark coordinates, and CT-to-atlas transformations of the six reference marmosets. BMA 2.0 and the reference data are distributed under the CC BY 4.0 license. Other data are available from the corresponding authors upon reasonable request.

---

**Supplementary Table S1. Results of statistical tests.**

All tests were two-sided (*n* = 6 target marmosets). *P* values for comparisons with intensity-based registration were Bonferroni-corrected for the three comparisons within each error measure and transformation model, and those for comparisons with the original landmarks for the two raters. Euclidean, mean Euclidean error; AP, DV, and ML, mean absolute errors along the anteroposterior, dorsoventral, and mediolateral axes; –, not applicable or not shown in a figure.

## Notes

https://github.com/H-Yagishita/Marmoset_CT-Atlas_Mapper

https://doi.org/10.5281/zenodo.23683937

### Competing Interest Statement

The authors have declared no competing interest.

